# choros: correction of sequence-based biases for accurate quantification of ribosome profiling data

**DOI:** 10.1101/2023.02.21.529452

**Authors:** Amanda Mok, Robert Tunney, Gonzalo Benegas, Edward W. J. Wallace, Liana F. Lareau

## Abstract

Ribosome profiling quantifies translation genome-wide by sequencing ribosome-protected fragments, or footprints. Its single-codon resolution allows identification of translation regulation, such as ribosome stalls or pauses, on individual genes. However, enzyme preferences during library preparation lead to pervasive sequence artifacts that obscure translation dynamics. Widespread over- and under-representation of ribosome footprints can dominate local footprint densities and skew estimates of elongation rates by up to five fold. To address these biases and uncover true patterns of translation, we present `choros`, a computational method that models ribosome footprint distributions to provide bias-corrected footprint counts. `choros` uses negative binomial regression to accurately estimate two sets of parameters: (i) biological contributions from codon-specific translation elongation rates; and (ii) technical contributions from nuclease digestion and ligation efficiencies. We use these parameter estimates to generate bias correction factors that eliminate sequence artifacts. Applying `choros` to multiple ribosome profiling datasets, we are able to accurately quantify and attenuate ligation biases to provide more faithful measurements of ribosome distribution. We show that a pattern interpreted as pervasive ribosome pausing near the beginning of coding regions is likely to arise from technical biases. Incorporating `choros` into standard analysis pipelines will improve biological discovery from measurements of translation.

## Introduction

Quantification of gene expression has enabled insights into many aspects of biology, often relying on mRNA abundances as a proxy for protein levels [1, 2]. Ribosome profiling, or ribo-seq, goes beyond mRNA quantification to generate a global snapshot of protein production through deep sequencing of ribosome-protected mRNA fragments (‘footprints’) [3]. In this method, nuclease digestion exposes the 20-30 nucleotides protected by actively translating ribosomes, which are then converted into a library of DNA molecules suitable for next-generation sequencing. Mapping of these fragments reveals which transcripts, and which regions within transcripts, are under active translation and at what levels. Ribosome profiling has led to numerous discoveries about translation and its regulation, from the detection of novel open reading frames to the identification of mechanisms underlying translational control [4, 5].

The quantitative aspect of ribosome profiling has proved a rich source of information from which to study translation dynamics. Each footprint recovered in a ribosome profiling dataset reflects the position of an individual ribosome on a transcript, resulting in precise codon-level resolution of the distribution of ribosomes across a transcript. Ribosomes are distributed unevenly along individual transcripts, reflecting biologically significant phenomena such as sites of ribosome pausing or stalling and inherent differences in elongation rate over different sequences. Per-position ribosome abundances can be used to infer translation elongation rates: the slower the elongation rate, the longer the dwell time and the more likely a ribosome is observed at that codon position. Analyses of ribosome profiling datasets have revealed that the largest driver of elongation rate is the identity of the codon being decoded in the aminoacyl (A) site of the ribosome, with relationships between elongation rate and tRNA availability, wobble base-pairing, and amino acid properties [6–10]. Elongation rate is also slightly impacted by the identity of the previous codon to have been decoded, now in the peptidyl (P) site along with its cognate tRNA, and perhaps by the identity of the tRNA being released in the exit (E) site. Mathematical and data-driven models of ribosome densities, fit from footprint counts, can capture the effects of synonymous codon choice to inform the design of transcripts for optimized translation efficiency [10–13]. Beyond these effects of intrinsic interactions and tRNA abundance, ribosome stalls and pauses can be caused by interactions of the ribosome with mRNA secondary structures, nascent peptide chains, or proteins involved in translation regulation, and these effects can also be seen as peaks of ribosome footprint density. For instance, ribosome profiling analysis has revealed sites of ribosome pausing due to binding of the fragile X mental retardation protein (FMRP), which mediates localized translation at synapses and contributes to synaptic plasticity [14]. The codon-level resolution of ribosome profiling therefore makes it a powerful method for discovering regulatory processes shaping gene expression.

However, experimental artifacts can distort estimates of translation rates, models of translation dynamics, and signals of ribosome pausing. A particularly strong distortion arises from sequence-based biases introduced during certain steps of library preparation protocols, namely, ligation of an adapter to the 3′ end of the footprint and circularization of the reverse transcription product in order to ligate an adapter to the 5′ end (Figure 1A). These biases result in over- and under-representation of certain footprints based on the nucleotide content at the fragment ends, leading to spurious sequence dependencies [13, 15, 16]. Similar sequence biases can arise from differences in ligation efficiency on structured RNA fragments [17, 18]. Sequence biases can even overshadow the impact of the codon being decoded by the ribosome, dramatically skewing footprint counts by up to five fold for purely technical reasons and severely limiting the insights that can be drawn from ribosome profiling data on individual genes.

**Figure 1.**
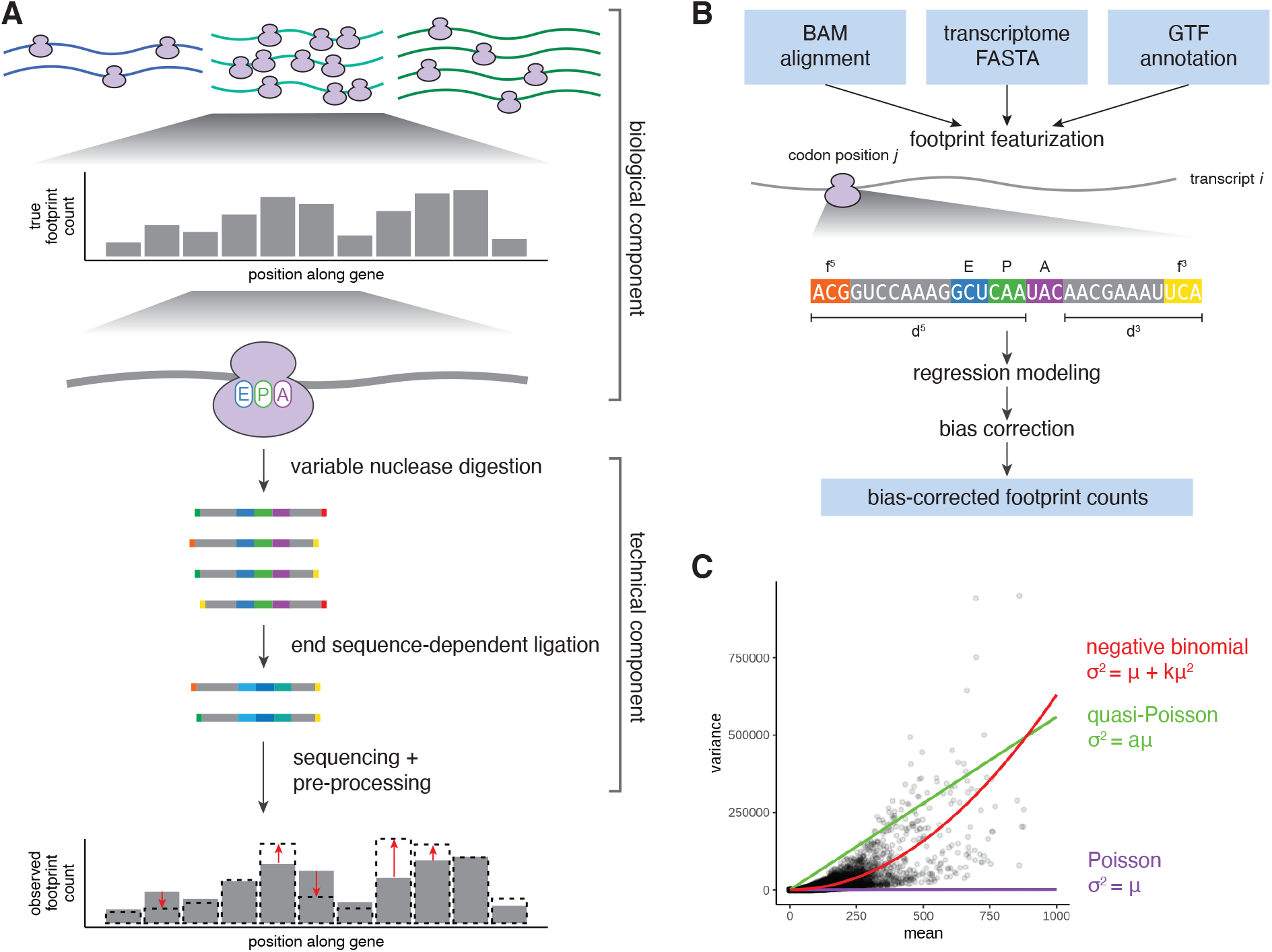
Overview of data-generating processes and bias correction with `choros`. **(A)** Ribosomes are distributed among transcripts based on transcript-specific mRNA abundances and translation initiation rates. Within a transcript, ribosome accumulation per codon position is related to the codons present in the ribosome A, P, and E sites, which largely determine the speed of elongation. Conditioned upon an individual ribosome, nuclease digestion occurs to differing levels of completion which leads to different nucleotides exposed at each end of the footprint sequence. Ligase enzymes in the library preparation protocol act upon these end sequences with different efficiencies, leading to differential recovery of footprint sequences depending on the nucleotides present at the footprint ends. **(B)** `choros` takes three input files: a BAM file of footprint alignments, a FASTA file of transcript sequences (including 5′ and 3′ UTRs), and a GTF annotation file specifying 5′ UTR, CDS, and 3′ UTR lengths. `choros` will then featurize each footprint alignment, aggregate footprint counts by transcript, codon position, and footprint length and 5′ frame, and compute a negative binomial regression model. Bias correction factors are adapted from regression coefficients and used to generate bias-corrected footprint counts. **(C)** Footprint counts are best modeled by a negative binomial distribution. Mean and variance estimates of per-codon-position footprint counts were calculated from a dataset generated with 14 wild-type replicates [23]. Mean-variance relationships corresponding to the Poisson, quasi-Poisson, and negative binomial distributions are shown in purple, green, and red, respectively.

Sequence preferences have been previously identified in other types of RNA sequencing experiments; however, in these cases the reads are summed across a full transcript or a wide window of sequence, and biases are likely to average out across the many different sequence contexts in one transcript [17–22]. When exploiting the codon-level resolution of ribosome profiling data to show variation in translation speed along a single transcript, the footprints at each position must be counted separately rather than summed. An explicit correction for sequence biases would improve the fidelity of ribosome profiling measurements. Slight variations in experimental protocols—for instance, differences in incubation time in ligation or digestion reactions—mean that the biases must be learned anew for each experiment. Direct experimental measurement of biases—for instance, by evaluation of an accompanying library generated in parallel from fractionated mRNAs—would be costly and add its own distortions [15]. Further, the over- or under-representation of different end sequences cannot be counted directly due to the confounding effects of the three-nucleotide periodicity in coding regions. To date, no computational method explicitly models the sequence-dependent biases to provide corrected ribosome footprint counts, but the rich data generated from ribosome profiling should render computational modeling quite feasible. The extensive sequence variation across millions of bases of sequence data is enough to reduce collinearity and make parameters of individual processes identifiable.

To this end, we present `choros`, a computational method that estimates biological and technical contributions to ribosome distributions and generates sequence bias-corrected footprint counts. We validate `choros` on data simulated with realistic sequence biases and demonstrate the method’s ability to capture underlying parameters. Applied to experimental ribosomal profiling datasets, `choros` correctly attenuates ligation biases and provides accurate estimates of relative ligation probabilities and codon-level ribosome occupancies. The extensible regression framework also enables detection of differential codon-level translation between experimental conditions. Using `choros`, we show that bias correction removes large artifacts that have been interpreted as biological signals and enables detection of biological phenomena such as ribosome collisions. `choros` is implemented as an open-source R package that can be incorporated into standard ribosome profiling analysis pipelines to provide accurate ribosome footprint counts for downstream analysis and biological discovery.

## Methods

Our software, `choros`, is provided as an R package for correcting sequence biases in ribosome profiling datasets and is available on GitHub at https://github.com/lareaulab/choros. simRiboSeq is an R package for simulating ribosome profiling data, available on GitHub at https://github.com/lareaulab/simRiboSeq. Code to regenerate figures in this work are available on GitHub at https://github.com/lareaulab/choros_paper.

### Regression modeling of biological and technical determinants of ribosome profiling datasets

Our method begins with a model of ribosome profiling data that is based on the physical processes that generate ribosome footprints: (i) a biological component that governs a true distribution of ribosomes, from which (ii) experimental protocols sample with bias to generate a sequencing library (Figure 1A). Overall ribosome abundances per gene are driven by mRNA abundances as well as transcript-specific translation initiation rates. Ribosome distributions along individual transcripts are driven by the speed of the ribosome as it decodes each codon position and translocates to the next position. The time required for this process depends on the identities of the codons in the ribosome A, P, and E sites, modeled by codon-specific terms for each site [13, 16, 24]. Nuclease digestion cleaves at variable positions around the edges of the ribosome, determining the total footprint length and the position of the 5′ and 3′ footprint ends relative to the codon being decoded in the A site (5′ and 3′ ‘digestion lengths’). This variable digestion exposes different nucleotides (‘bias sequence’) at the footprint 5′ and 3′ ends, which are ligated at different efficiencies due to sequence preferences of the enzymes used in library preparation protocols. Footprint recovery may additionally be impacted by secondary structure of the RNA fragment, which we model with GC content of the footprint excluding A, P, and E site codons [17].

We fit a generalized linear model (GLM) to estimate the parameters underlying this data-generating process (Equation 1). The model assumes that the number of footprints 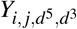 mapped to a transcript *i* at codon position *j* with 5′ and 3′ digestion lengths *d*^5^ and *d*^3^, respectively, follows a negative binomial distribution with mean 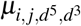 and dispersion parameter *φ*. The negative binomial distribution allows observed extra variability that cannot be accounted for by Poisson or quasi-Poisson regression models for count data. We denote the codons residing in the footprint’s A, P, and E sites as *A*_*i, j*_, *P*_*i, j*_, and *E*_*i, j*_, respectively. The bias sequences at the 5′ and 3′ ends of the footprint are denoted 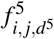 and 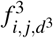 respectively. The footprint’s overall GC content is denoted as 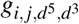 and encoded as a continuous covariate. We can then model the observed footprint counts as:

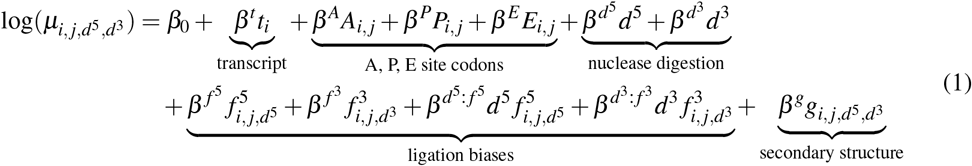

where

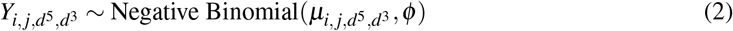

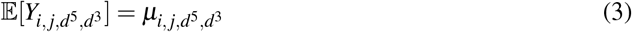

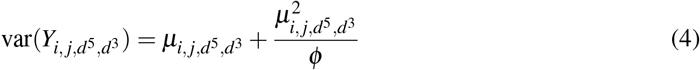

We include interaction terms for digestion length and bias sequence at each end of the footprint to capture possible sequence preferences of nuclease digestion as well as effects of non-templated nucleotide additions during reverse transcription [25]. The regression framework is extensible and can be amended to include additional terms, such as an interaction term to estimate differential A-site codon effect between experimental conditions (Equation 5):

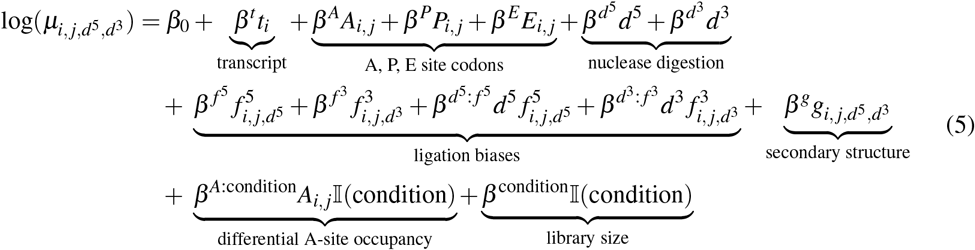

To reverse the effects of biased footprint recovery, we generate a correction factor for each individual footprint based on the parameter estimates learned in the regression fit (Equation 6):

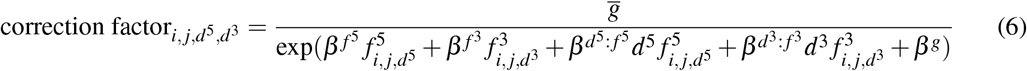

where 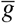 is the mean GC content of the observed footprint reads. Read counts are then multiplied by these correction factors and scaled to maintain total library size in order to produce bias-corrected counts.

### Overview of choros

`choros` has four major modules: data processing (including A-site assignment and bias sequence determination), regression fitting, bias correction, and bias evaluation (Figure 1B). The inputs to `choros` are as follows: alignments of ribosome profiling reads (i.e. BAM file), a reference transcriptome (i.e. FASTA file), and a gene annotation file that defines 5′ UTR, CDS, and 3′ UTR regions within the reference transcriptome. After fitting the model with `glm.nb` from the MASS package and performing bias correction, `choros` outputs a table of raw and corrected read counts, aggregated by transcript, codon position, and 5′ and 3′ digestion lengths.

Footprints were assigned to transcript codon positions using an A-site assignment process previously described in Tunney et al. [13]. In brief, A-site offset rules were determined per footprint length and frame of the 5′ terminal nucleotide relative to the coding sequence, based on a metagene plot of alignments near the start codon (Supplementary Figure 1). The canonical 28-nt footprint in frame 0 is assigned an offset of 15, indicating that the A-site codon starts in the 16th nucleotide position of the read. Offset rules for other lengths and frames were chosen to be consistent with over- and under-digestion relative to the 28-nt footprint. We then took the first two or three nucleotides of the footprint as the 5′ bias sequence and the last three nucleotides as the 3′ bias sequence. All sequences were pulled from the underlying transcript sequence to avoid sequencing errors and mismatches in the read itself.

`choros` is flexibly designed to fit the regression model on a user-supplied list of transcripts and codon positions. While the technical biases that arise during library preparation should be universal to footprints from all transcripts, sparsity in read coverage may lead to unreliable parameter estimates. Unless otherwise described, all regression models in this work were fit to a random set of 250 of the top 500 transcripts according to footprint coverage (mean footprint abundance per codon position) after excluding transcripts that had fewer than 5 mean reads per codon position and fewer than 100 codon positions with mapped footprints. In addition, the first and last 20 codon positions of transcripts were excluded from the regression fit to avoid the effects of translation initiation and termination.

Bias correction was performed as described above (Equation 6). Biases before and after correction were evaluated using a method similar to the feature importance measurements as described in Tunney et al. [13]. In brief, per-codon-position footprint counts in truncated coding sequences were scaled by the average footprint count across the coding sequence to control for transcript-specific mRNA abundances and translation initiation rates. Input features were defined as a neighborhood of 13 codons (or corresponding number of nucleotide positions) centered around that codon position. A series of linear regression models predicting scaled footprint count from the input sequence neighborhood were fit, where the full model included the entire sequence neighborhood as input features and subsequent leave-one-out models omitted one codon or nucleotide position at a time. The feature importance of each position was computed as the difference in correlation of the true and predicted footprint counts between the full and corresponding leave-one-out model. Position importance plots generated from these feature importance metrics were used to evaluate for the presence of recovery biases.

### Data simulation

To test `choros`, we developed an R package `simRiboSeq` to simulate ribosome profiling datasets with realistic parameters for individual steps in the data-generating process (Figure 1A, Supplementary Figure 2). Per-transcript footprint counts are generated by multinomial sampling with user-defined transcript probabilities. Given a transcript sequence and ribosome count, per-codon-position footprint counts are generated by multinomial sampling where the probability per codon position is proportional to a user-defined codon-specific value. Each footprint count generates an individual footprint sequence, where the 5′ and 3′ digestion lengths are sampled with user-defined probabilities. Whether footprint sequences are retained is simulated as a Bernoulli process, where the probabilities of success are defined by the 5′ and 3′ bias sequences separately. These steps are performed iteratively until the desired library size has been achieved. Footprint sequences are written to file in FASTQ format.

Values for simulation probabilities were adapted from experimental datasets. Per-transcript probabilities were generated from per-transcript ribosome abundances extrapolated from a loess regression of per-transcript ribosome abundances on transcript lengths fit to data from Weinberg et al. with additional Poisson noise [26]. Per-codon values were adapted from the A-site codon weights from a neural network model of ribosome distributions trained on data generated by Weinberg et al. [13, 26]. 5′ and 3′ recovery probabilities were adapted from the *−*5 and +3 codon weights from a neural network model of ribosome distributions trained on data generated by Schuller et al. [13, 27].

### Pre-processing of ribosome profiling datasets

The experimental datasets analyzed in this work are listed in Table 1 [13, 23, 26–30]. Reference ribosomal RNA, noncoding RNA, and annotated coding sequence files were previously described, with the modification that ORF sequences include 20 nt of 5′ UTR and 20 nt of 3′-UTR sequences to accommodate all footprint reads at start and stop codons [13].

**Table 1.**
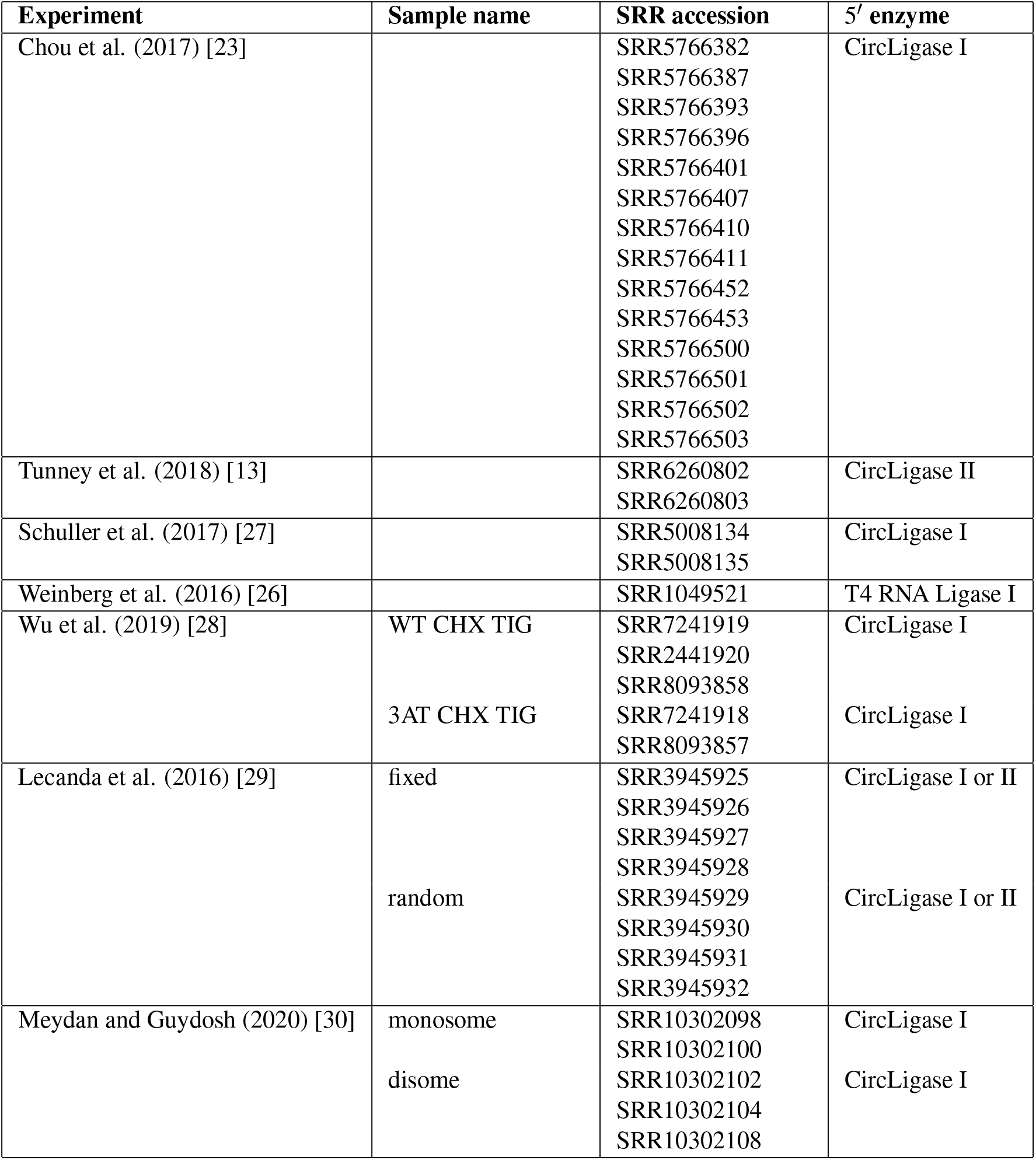
Experimental ribosome profiling datasets analyzed Weinberg et al. used T4 RNA Ligase 1 for 3′ adapter ligation; all other datasets used T4 RNA Ligase 2 K227A.

Raw ribosome profiling reads were downloaded from the NCBI Sequence Read Archive using fasterqdump (SRA Toolkit, https://github.com/ncbi/sra-tools). Trimming of 3′ adapters was performed with cutadapt, and trimmed reads longer than 20 nt were retained with options ‘--trimmed-only -m 20’ [31]. If the 3′ adapter or the reverse-transcription (RT) primer contained random nucleotides, those were extracted as unique molecular identifiers (UMIs) for deduplication with umi tools extract [32]. Reads were aligned, allowing two mismatches, to ribosome and noncoding RNAs using bowtie v.1.0.0 (chosen due to the ability to explicitly specify maximum allowable mismatches) with the options ‘-v 2 -S --un’, and unaligned reads were kept for downstream processing [33].

If UMIs were extracted, deduplication was performed with the following steps. Since UMI-tools deduplicates reads by alignment coordinate and read length, we aimed to provide the tool a SAM alignment file with only one alignment reported per read. However, in the case of multi-mapping reads, reads derived from the same original molecule (i.e. containing the same UMI) may not necessarily be assigned the same alignment locus and therefore would not be properly deduplicated. To control for this scenario, we instructed bowtie to report all best alignments and then manually selected the positionally first alignment per sequencing read. Thus, the following procedure was developed. First, reads were aligned to the transcriptome using bowtie, allowing two mismatches, and all best alignments were reported with the options ‘-v 2 -S --norc -a --best --strata’. The resulting alignment file was then sorted by read name using samtools sort with options ‘-n -O SAM’ [34]. Per read, the positionally first alignment was retained and remaining alignments were discarded using a custom R script. The alignment file was then sorted by alignment coordinate with samtools sort and indexed with samtools index in preparation for deduplication. Deduplication was performed using UMI-tools using read length in addition to alignment coordinates and UMI to identify duplicates, with options ‘--read-length’. Unique reads were then converted into FASTQ format with samtools fastq.

Finally, we aligned the remaining reads to the transcriptome to quantify translation. Ribosome footprints may not map uniquely to a single transcript due to the short length of the sequencing read and the presence of multiple transcript isoforms or gene paralogs with high sequence similarity. To address this, we report all valid alignments and use RSEM to assign each alignment a posterior probability that we interpret as a fractional read count [35]. De-duplicated, non-rRNA reads were aligned to the transcriptome using bowtie, allowing all alignments with options ‘-v 2 -S --norc -a -l 15’. Mapping weights for multi-mapping reads were computed using RSEM with options ‘--seed-length 15’. After A-site assignment, for each codon position and combination of 5′ and 3′ digestion lengths, raw footprint counts were computed by summing the RSEM mapping weights for each corresponding alignment.

## Results

### choros accurately estimates ligation biases in simulated data

The negative binomial distribution is conventionally used to model count data arising from RNA-seq experiments because it can account for the observed extra-Poisson variability. We first wanted to confirm the appropriateness of negative binomial regression for ribosome profiling data, in keeping with previous applications [36–39]. Using per-codon-position ribosome footprint counts from a published dataset of 12 yeast ribosome profiling replicates [23], we found that a negative binomial distribution with the variance as a quadratic function of the mean fit the data well, while a Poisson or quasi-Poisson distribution did not capture the observed variability (Figure 1C).

We next evaluated `choros` on simulated data to test how well the regression framework quantified the parameters used to generate the data and how well the computed correction factors were able to attenuate footprint end sequence biases. Data were simulated with `simRiboSeq` where per-codon A-site parameters and biased ligation parameters were adapted from weights learned by models reported in Tunney et al. [13]. Four simulated datasets were generated to evaluate the following scenarios: (i) footprint ligation is not biased; (ii) ligation of the 3′ adapter is affected by the footprint 3′ bias sequence; (iii) ligation at the footprint 5′ end is affected by the footprint 5′ bias sequence; and (iv) footprint recovery is impacted by both the 3′ and 5′ bias sequences.

We fit the `choros` regression model to these simulated data and evaluated how well the learned regression coefficients reflected simulation parameters. Of particular interest were the 5′ and 3′ ligation bias terms, as they are used to generate bias correction factors. Across all four simulated datasets, the regression coefficients precisely recapitulated simulation parameters (Figure 2A). The 5′ and 3′ ligation bias terms were assigned the appropriate values corresponding to their simulation probabilities and took on null values where no ligation biases were simulated. The A-site regression coefficient terms mirrored codon-level probabilities, supporting their utility in inferring codon-specific translation elongation rates in experimental data.

**Figure 2.**
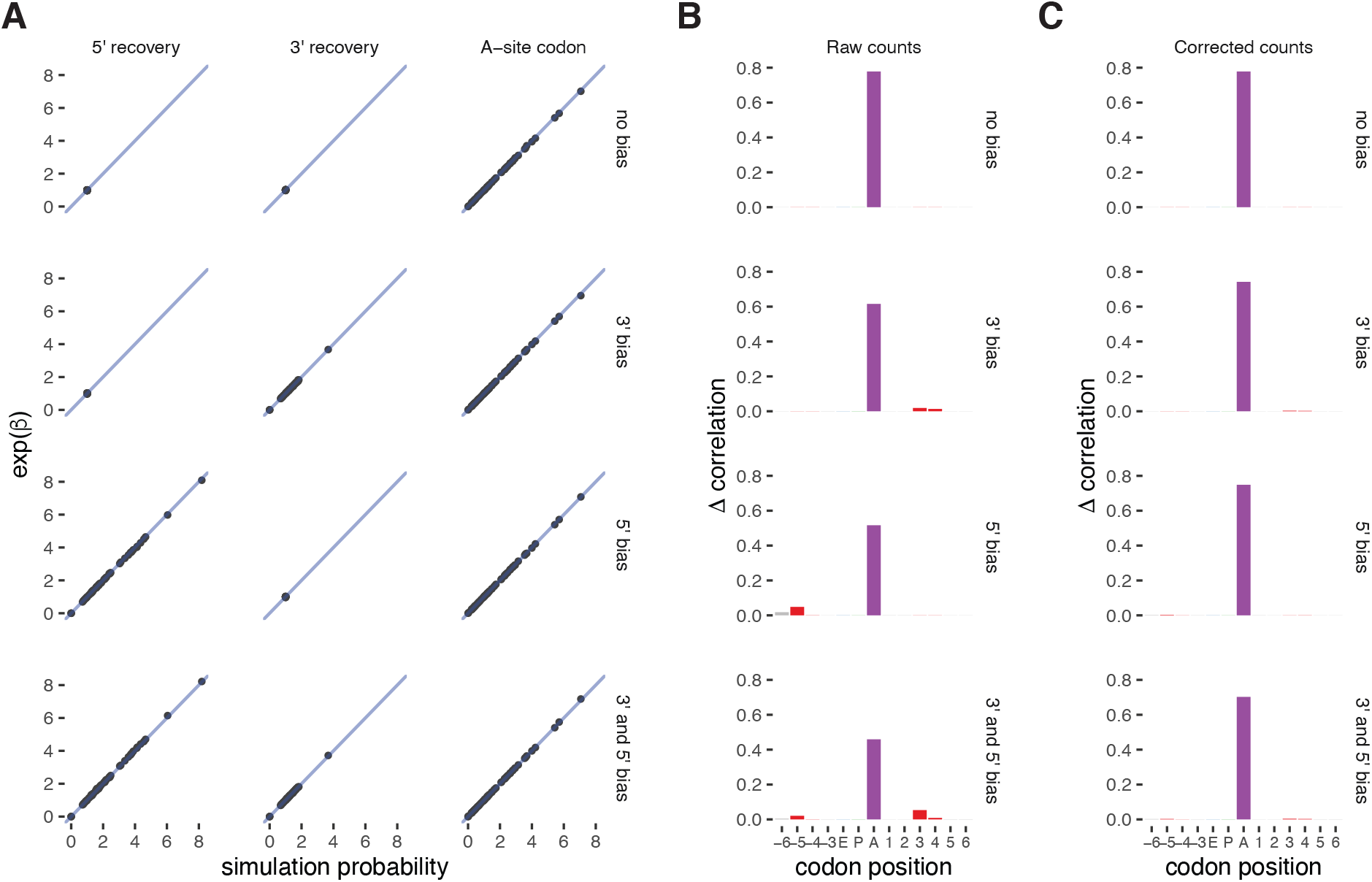
`choros` correctly models theoretical data-generating process. **(A)** Comparison of simulation parameters used by `simRiboSeq` to generate synthetic ribosome profiling data and regression coefficient estimates learned by `choros`. Simulation parameters are normalized to the corresponding reference sequence and regression coefficients are exponentiated for proper comparison. Each point represents one three-nucleotide sequence at the footprint 5′ or 3′ end or in the ribosome A site. The 1:1 line is shown in blue. **(B)** Position importance plots generated for simulated data. The height indicates the improvement in correlation between observed and predicted footprint counts for a full regression model versus one in which the features at a given position is left out of the model. **(C)** Position importance plots for bias-corrected simulated data.

To test whether bias correction factors could attenuate the sequence-based signals at the ribosome boundaries, we applied the bias correction procedure to simulated footprint counts and evaluated both the raw and corrected counts for the presence of ligation biases. The position importance plots generated for raw counts demonstrated position contributions only where expected, namely at the ribosome A site, at the *−*5 and *−*6 codon positions (corresponding to biased 5′ ligation), and at the +3 and +4 codon positions (corresponding to biased 3′ ligation) (Figure 2B). Before correction, the datasets with simulated ligation biases show a decreased A-site signal relative to the ‘no bias’ dataset, showing that biased recovery obscures biological signal. After correction, the signal at the footprint boundaries (positions *−*6, *−*5, +3, and +4) is properly lowered to negligible levels for the datasets with simulated ligation biases, and the A-site contribution increases to match the ‘no bias’ datasets (Figure 2C). No change in position importance was observed in the ‘no bias’ dataset after bias correction, indicating that the bias correction procedure does not improperly adjust footprint counts when ligation bias is not present. We also note that the lack of signal at other codon positions after correction suggests that the bias correction process does not cause distortions due to properties of natural transcript sequences such as correlations between adjacent codons. Taken together, `choros` is able to generate bias correction factors that reverse the effects of ligation biases in simulated ribosome profiling data.

### Bias correction attenuates sequence artifacts in ribosome profiling datasets

Ribosome profiling protocols vary across datasets, from the use of different ligases and adapter sequences to differences in experimental conditions. The overall structure of the model of ribosome footprint generation remains the same, with the variation between datasets arising from differing contributions of the components in the data-generating process (e.g., biological differences in decoding speed due to differences in tRNA abundance; technical differences in sequence bias due to varying preferences of different enzymes). `choros` should therefore be trained separately per dataset to capture these differences and generate bias correction factors specific to each dataset.

We evaluated `choros` on three ribosome profiling datasets generated in the budding yeast *Saccharomyces cerevisiae* [13, 26, 27]. We observed strong contributions of the sequence in the ribosome A site, expected to correspond to tRNA decoding, as well as at the −5 and +3 codon positions, expected to correspond to sequence biases (Figure 3A). After bias correction, the *−*5 and +3 position signals decreased while the A-site signal increased, indicating that the corrected counts show much less impact of ligation biases and more strongly reflect true ribosome distributions (Figure 3B). As expected, the A-site coefficients estimated by `choros` correlated with the tRNA adaptation index (tAI), a widely used measure of codon preference, across all three datasets (Figure 3C). Less adaptive codons should be translated more slowly and thus correspond to higher ribosome occupancies, leading to the observed negative correlation. Importantly, `choros` was able to recover the signatures of known sequence bias. For the datasets that used CircLigase in the 5′ ligation step, the 5′ bias regression coefficients estimated by `choros` correlated strongly with our previous *in vitro* measurements of CircLigase ligation efficiencies (Figure 3D) [13]. As expected, the 5′ bias coefficients calculated for a dataset that used T4 RNA ligase 1 rather than CircLigase showed no correlation with CircLigase efficiencies.

**Figure 3.**
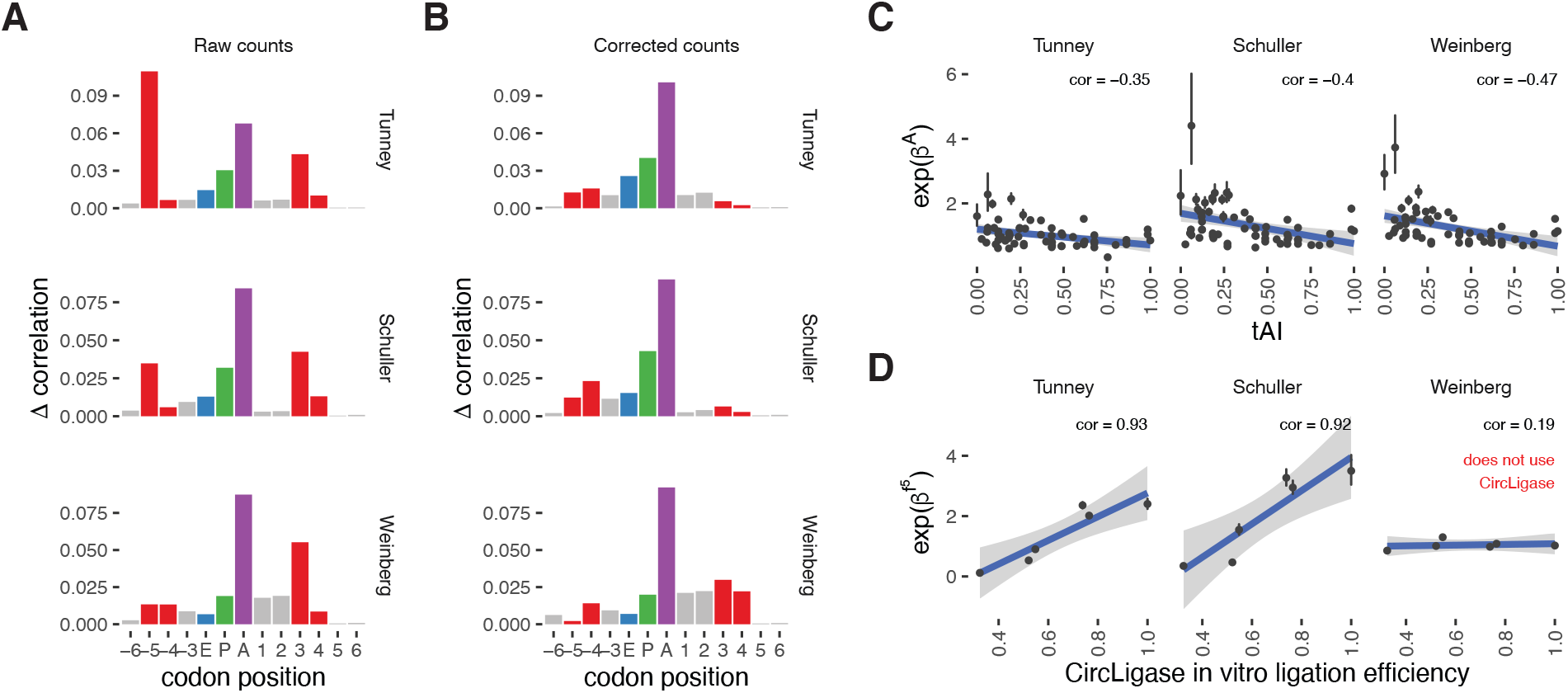
`choros` attenuates ligation biases and learns real-world parameters in yeast ribosome profiling datasets. **(A)** Position importance plots for raw footprint counts in three yeast ribosome profiling datasets. Red denotes the codon positions corresponding to the ribosome boundary that are assumed to be affected by technical, rather than biological, factors. **(B)** Position importance plots for bias-corrected footprint counts. **(C)** `choros` regression coefficients for A-site codon occupancies capture translation elongation dynamics as shown by their negative correlation with tAI. **(D)** `choros` regression coefficients match experimental measurements of *in vitro* ligation efficiency (data from Tunney et al. [13]). The experiments that used CircLigase in experimental protocols had 5′ bias regression coefficient estimates that correlated strongly with CircLigase ligation efficiencies. The Weinberg data did not use CircLigase, and those coefficients did not correlate with CircLigase efficiency measurements.

We found that our bias correction approach shifted the distribution of ribosome footprint counts over the individual positions of a transcript, but with no major effect on total footprints per transcript (Supplementary Figure 3A-B). This suggests that the biased ligation of different footprint end sequences is largely averaged out across the diversity of footprint sequences originating from one gene. This averaging effect is also apparent in codon pause scores, where footprint abundances are aggregated over all occurrences of a particular codon and therefore also reflect a diversity of footprint end sequences (Supplementary Figure 3C). Thus, while technical biases in ribosome profiling could have a small effect on overall measurements of translation or estimates of the relative decoding speed of synonymous codons, the key contribution of `choros` is less-distorted estimates of the distribution of ribosomes along transcripts from a single gene, enabling codon-level resolution for discovery of translation dynamics.

### Footprint counts are more concordant after bias correction

To validate the ability of `choros` to recover estimates of ribosome distribution that are closer to a ground truth, we applied our method to ribosome profiling data from an experiment in which material was split and subjected to different protocols for library generation. In this scenario, the underlying distribution of ribosomes should be consistent between the ribosome profiling datasets, and divergences between the datasets would be driven by disparate biases from distinct library preparation protocols. Removing these artifacts with a bias correction procedure should then recover consistency between those datasets. To test this idea, we applied `choros` to datasets generated by Lecanda et al. where a single batch of yeast lysate was digested, footprint fragments were extracted, and then the purified footprint fragments were split into two batches that were each subjected to a different protocol: (i) 3′ adapter ligation, reverse transcription, and circularization with no randomized sequences (‘fixed adapter’); and (ii) a dual-randomization strategy (‘random adapter’) using a 3′ DNA adapter with 4 nt of randomized nucleotides at its 5′ end and an RT primer with 3 nt of randomized nucleotides at its 5′ end (Figure 4A) [29]. While the ligase would encounter the same footprint end sequences in either case, preferences for the end nucleotides on the linker also affect ligation efficiencies, and the ligase may prefer certain pairings of footprint and linker end sequences [22]. We therefore expect the ligation biases to differ between the fixed-adapter and random-adapter datasets, and use these datasets as a test case to evaluate concordance after bias correction with `choros`.

**Figure 4.**
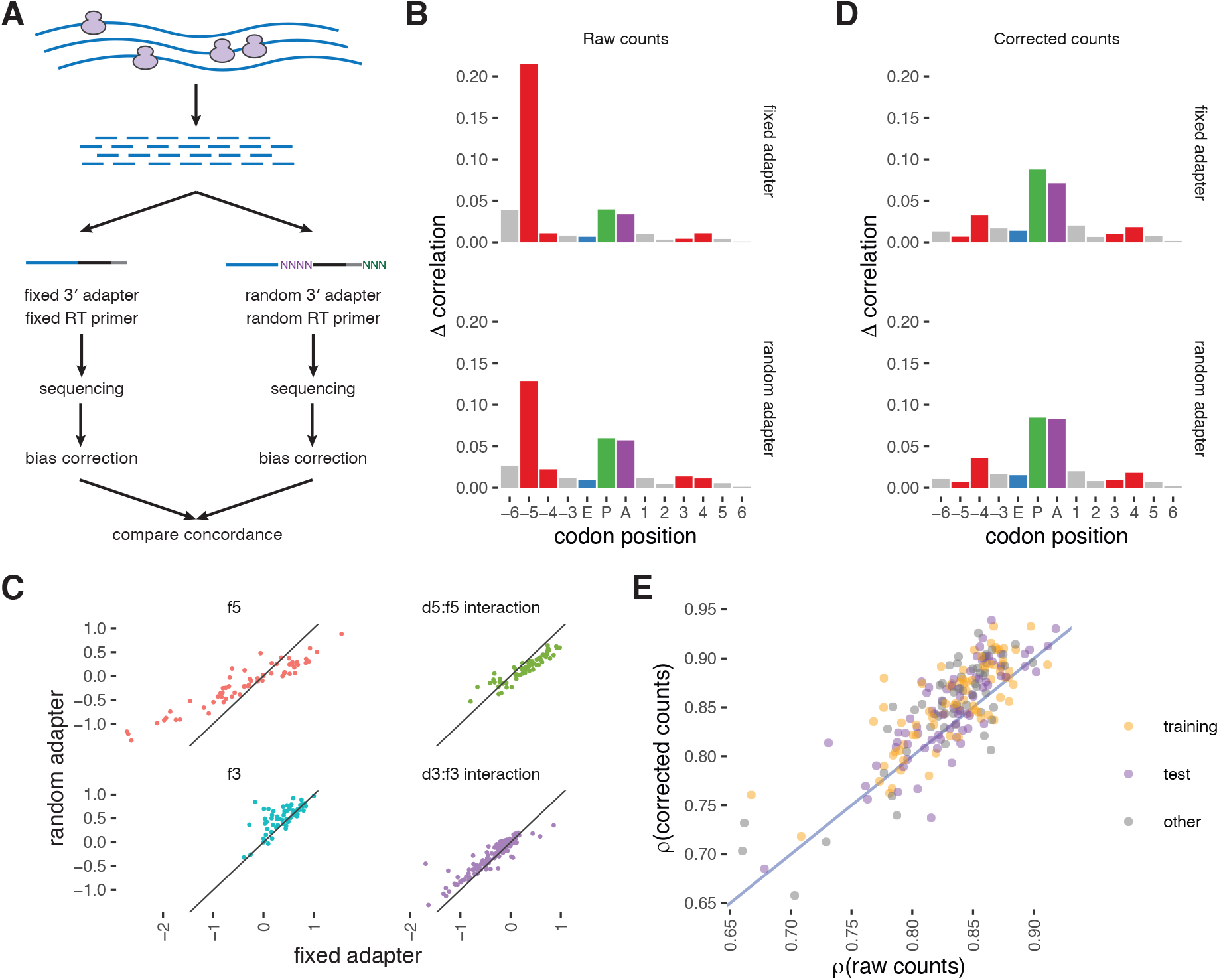
Direct comparison of bias-corrected footprint counts demonstrates higher concordance on well-translated genes. **(A)** Schematic for the generation and analysis of an experimental dataset generated by Lecanda et al. for validation of `choros` [29]. A pool of ribosome footprints was split to undergo separate library preparation protocols: a standard protocol and (ii) a protocol that involved ligation of the footprint 3′ and 5′ ends onto adapters ending with several random nucleotides. These libraries separately underwent next-generation sequencing and bias correction with `choros`. We compare concordance of the datasets before and after bias correction. **(B)** Position importance plots for raw footprint counts. Substantial signal at the ribosome 5′ boundary is observed. Use of an RT primer with random nucleotides roughly halved the sequence-dependent biases at the footprint 5′ end. **(C)** Comparison of regression coefficients between datasets. **(D)** Position importance plots for bias-corrected footprint counts. **(E)** Evaluation of dataset concordance before and after bias correction. Each point represents a single transcript with average coverage of at least 50 footprints per codon position. Spearman correlation of raw footprint counts per position between the fixed-adapter and random-adapter datasets was compared to Spearman correlation of bias-corrected counts between the two datasets.

In keeping with prior results [25, 29], we found that randomization of the 5′ end of the RT primer, which is ligated by CircLigase to the 3′ end of the reverse-transcribed cDNA (corresponding to the 5′ end of the original footprint), significantly lessened the bias towards certain 5′ footprint sequences. This effect can be seen as a lesser, albeit still high, contribution of the 5′-most codon position in the position importance plots for the randomized adapter dataset (Figure 4B). In contrast, randomization of the 5′ end of the adapter ligated to the 3′ end of the original footprint had a relatively small impact on overall footprint counts. This effect was reflected in `choros` regression coefficients. The 5′ bias parameter estimates for the random-adapter dataset were dampened relative to the fixed-adapter dataset (Figure 4C), while 3′ bias parameter estimates only varied slightly between the two datasets. Regression coefficients for A-, P-, and E-site codons correlated strongly between datasets, consistent with the fact that the libraries were generated from the same pool of ribosome footprints (Supplementary Figure 4).

We then confirmed the ability of `choros` to attenuate these biases. After bias correction, the position importance plots showed a decrease in signal at the ribosome boundary positions and an increase in signal at the ribosome P and A sites (Figure 4D). The plots for the two datasets became remarkably similar, indicating that the differences observed in the raw counts were driven by ligation biases that were correctly reversed by `choros`. These results indicate that the inclusion of random nucleotides in the RT primer is an effective method to attenuate some but not all ligation biases, and that `choros` is able to correct the remaining sequence artifacts present in these datasets.

Next, to see if bias correction enabled more accurate measurement of footprint distribution along individual transcripts, we evaluated the concordance in footprint distribution between the fixed-adapter and random-adapter datasets, before versus after bias correction. Footprints were aggregated by A-site codon position, and footprint counts were compared between the two datasets. The two datasets were indeed more consistent after bias correction, with the overall Spearman correlation between datasets increasing from 0.73 to 0.84. The improvement from bias correction varied by transcript, with a strong dependence on transcript coverage (Supplementary Figure 5). While the transcripts with the highest average footprint coverage showed an increase in correlation after bias correction, transcripts with sparse footprint coverage actually became more discordant after bias correction. We found that bias correction improved the correspondence of footprint distributions on transcripts that had an average of 50 or more footprints per codon (Figure 4E). In all, our results emphasize that high coverage and low sparsity are critical both for generating reliable ribosome profiles and for estimating ligation biases. For the relatively small number of transcripts whose high coverage allows biological interpretation of codon-by-codon ribosome distribution along an individual transcript, bias correction can be applied successfully to improve estimates of ribosome dwell time on individual positions.

### Correction of sequence bias removes apparent ribosome pauses

Spikes in footprint counts—whether caused by ligation biases or genuine translational stalls or pauses— can be visible in ribosome profiling data on individual genes, but these signals are expected to be averaged out in aggregate data such as metagene plots showing the overall distribution of ribosomes over coding regions. However, some pervasive, position-specific spikes do appear in aggregate data. The overabundance of footprints on the sixth codon of reading frames, visible in metagene plots, has been interpreted as an indicator of a pause while the ribosome commits to elongation [40]. We reasoned that this overabundance could be caused by ligation bias rather than a mechanistic detail of translation. Most footprints from ribosomes decoding the sixth codon of a reading frame have their 5′ end 15 nucleotides upstream, exactly at the AUG start codon (Figure 5A). If fragments beginning with AUG are a particularly good substrate for ligation, these footprints will be recovered disproportionately.

**Figure 5.**
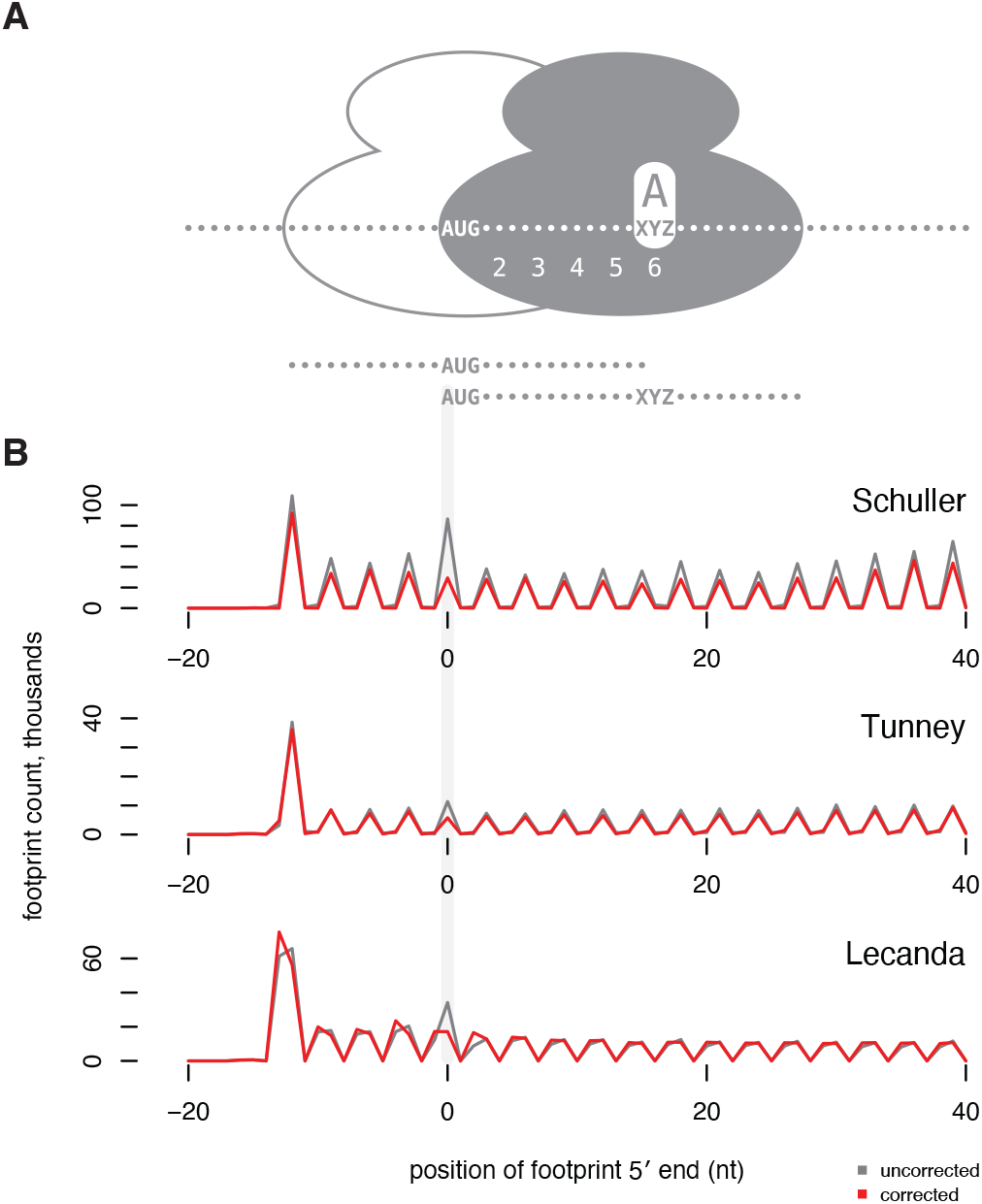
Signal of apparent ribosome pausing at the sixth codon of open reading frames is removed by bias correction. **(A)** A ribosome decoding the sixth codon of a transcript, with that codon in its A site, protects a footprint with its 5’ end at the start codon, as shown by the solid ribosome. For reference, the outlined ribosome shows the first ribosome on a reading frame, which begins with the AUG codon in its P site and decodes the second codon in its A site. The 28-nt fragments protected by these ribosomes are shown below. **(B)** Metagene plots show the count of footprints with a 5′ end mapping to each position relative to the start codon, summed across all genes. Position 0 shows footprints with their 5′ end aligned with the start codon, inferred to be decoding the sixth codon. Footprint counts before `choros` correction (grey) show a pronounced peak at the sixth codon that is absent after `choros` correction (red). Plots include the single most common footprint size from Schuller et al. (28 nt), Tunney et al. (28 nt), and Lecanda et al. (29 nt).

We observed a strong peak of ribosome footprints at the sixth codon in all of the analyzed datasets that were generated with CircLigase protocols (Figure 5B). After bias correction, this signal disappeared completely. To confirm that AUG was a preferred substrate for ligation, we inspected regression co-efficients and found that in all three datasets, AUG was among the top three 5′ regression coefficient estimates. We note that the codons near the beginning of transcripts are ignored in training the models, so this bias was recovered independently. Thus, computational inference and correction of technical biases can remove artifacts that have confounded biological interpretations.

### Detection of differential A-site ribosome occupancy

`choros` fits a generalized linear model to model the biological and technical processes underlying ribosome profiling data. The extensibility of this regression framework allows the inclusion of additional terms. One such parameter is the detection of differential codon-level ribosome occupancy under two experimental conditions (Equation 5). The decoding speed of a codon depends on factors including availability of its cognate tRNA, which can vary between cell types or in different metabolic conditions, adding a layer of control to translation. Codon-level occupancies or pause scores, which are empirically computed in various ways, lack statistical properties that allow detection of true versus spurious differences. In contrast, adding an interaction term to capture sample-specific A-site codon occupancy in the `choros` model simultaneously enables the statistical inference of differential codon-level occupancies that are independent of ligation biases. As quantification of tRNA species remains a technically difficult task due to complex secondary structures and diverse mRNA modifications, a statistical framework for differential codon occupancy may serve as an attractive proxy for detecting differences in tRNA abundances that drive differences in translation output [41, 42].

We assessed this approach on a dataset where yeast cells were treated with 3-amino-1,2,4-triazole (3-AT), a known inhibitor of histidine biosynthesis [28]. When fitting the `choros` model to each condition separately, the A-site regression coefficients for histidine codons, CAC and CAU, had substantially higher values in the sample treated with 3-AT, consistent with stronger ribosome pausing and slower decoding at histidine codons (Figure 6A). To evaluate whether these differences were statistically significant, we next fit footprint counts from both conditions simultaneously under a single `choros` regression model with an interaction term corresponding to A-site occupancy in the 3-AT condition. Footprints containing CAC and CAU codons in the A site were 41 and 70 times more enriched in the 3-AT sample relative to untreated, respectively (Figure 6B, p *<* 1e-25). Thus, `choros` is able to detect and quantify deviations in codon decoding speeds between experimental conditions.

**Figure 6.**
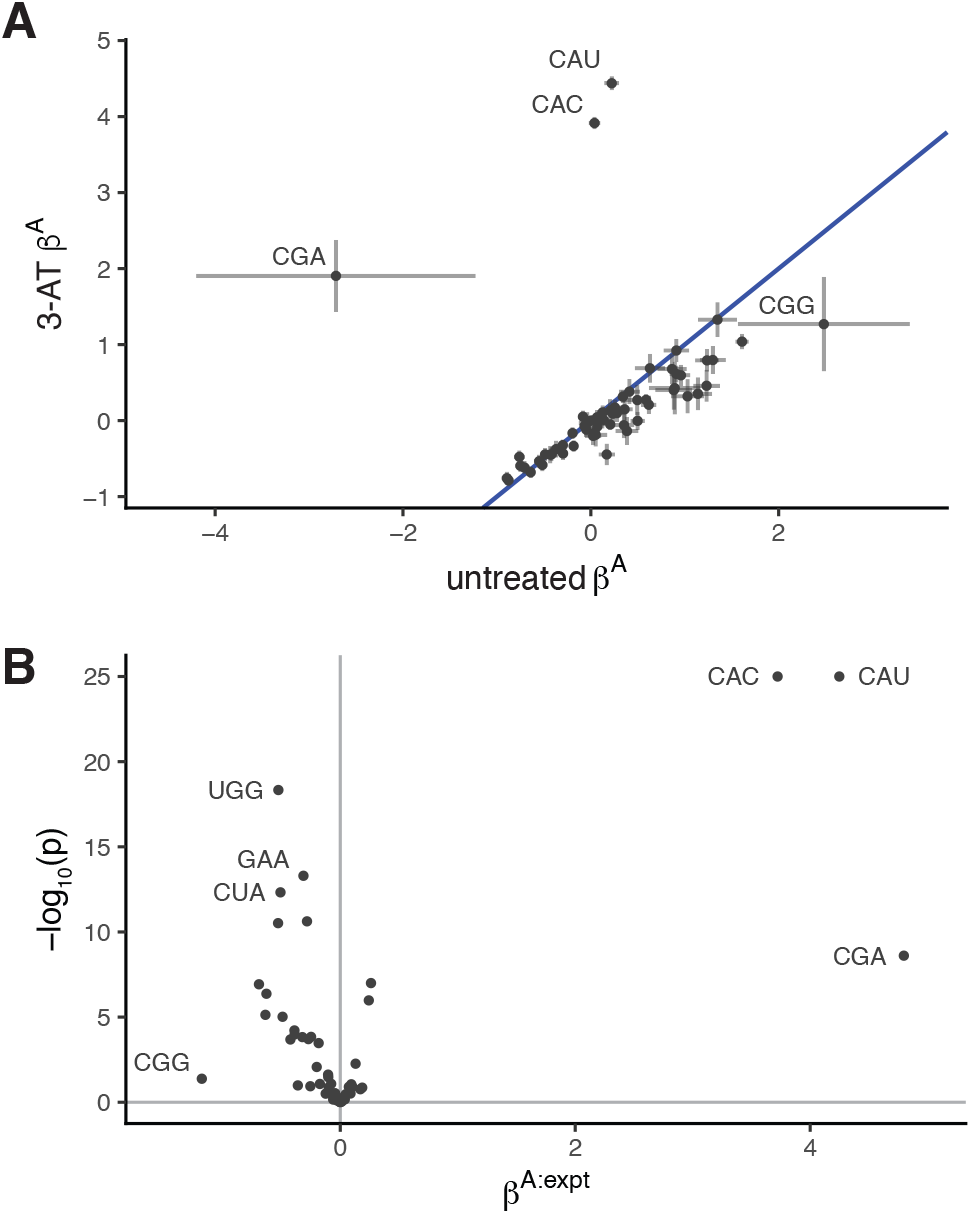
Detection of differential codon-level ribosome occupancy with `choros`. **(A)** `choros` regression coefficients capture higher ribosome occupancy at histidine codons (CAU and CAC) following treatment with 3-AT, an inhibitor of histidine biosynthesis. The pattern between untreated and 3-AT A-site regression coefficients (*β* ^A^) closely mimics the codon pause scores originally reported by Wu et al. [28]. Error bars represent the 95th percentile confidence interval of regression coefficient estimates; we note that there are very few CGA and CGG codons in yeast genes and their coefficient estimates are highly variable. **(B)** Volcano plot of differential ribosome A-site occupancy. `choros` parameters were estimated simultaneously for both datasets, incorporating an interaction term between A-site codon identity and experimental condition, *β* ^A:expt^, to capture differential A-site occupancies.

### Bias correction factors learned from monosome data attenuate biases in disome data

Ribosome collisions are widespread throughout the transcriptome, particularly on highly translated transcripts loaded with multiple ribosomes [30, 43–45]. Of much recent interest, disomes (two collided ribosomes) not only point to sites of endogenous ribosome pausing, but may also reveal the cellular processes involved in the resolution of ribosome stalls and the impact of collisions on protein output and cellular stress responses.

Collisions of a 5′ elongating ribosome into a 3′ paused ribosome can result in longer protected fragments, or disome footprints, that can be recovered by ribosome profiling using a broader size selection (Figure 7A). Analyses of these footprints may reveal dynamics underlying disome formation and collision resolution, but these footprint counts are subject to the same technical artifacts present in monosome datasets, obscuring the true sites of ribosome collisions. The more complicated biological process resulting in disomes, as well as the sparsity of disome footprints, makes modeling ligation biases difficult. However, analysis of the abundant monosome footprints generated in parallel to disome footprints can be leveraged to generate bias correction terms that can be applied to disome datasets, thus obtaining disome counts free of ligation-induced sequence artifacts.

**Figure 7.**
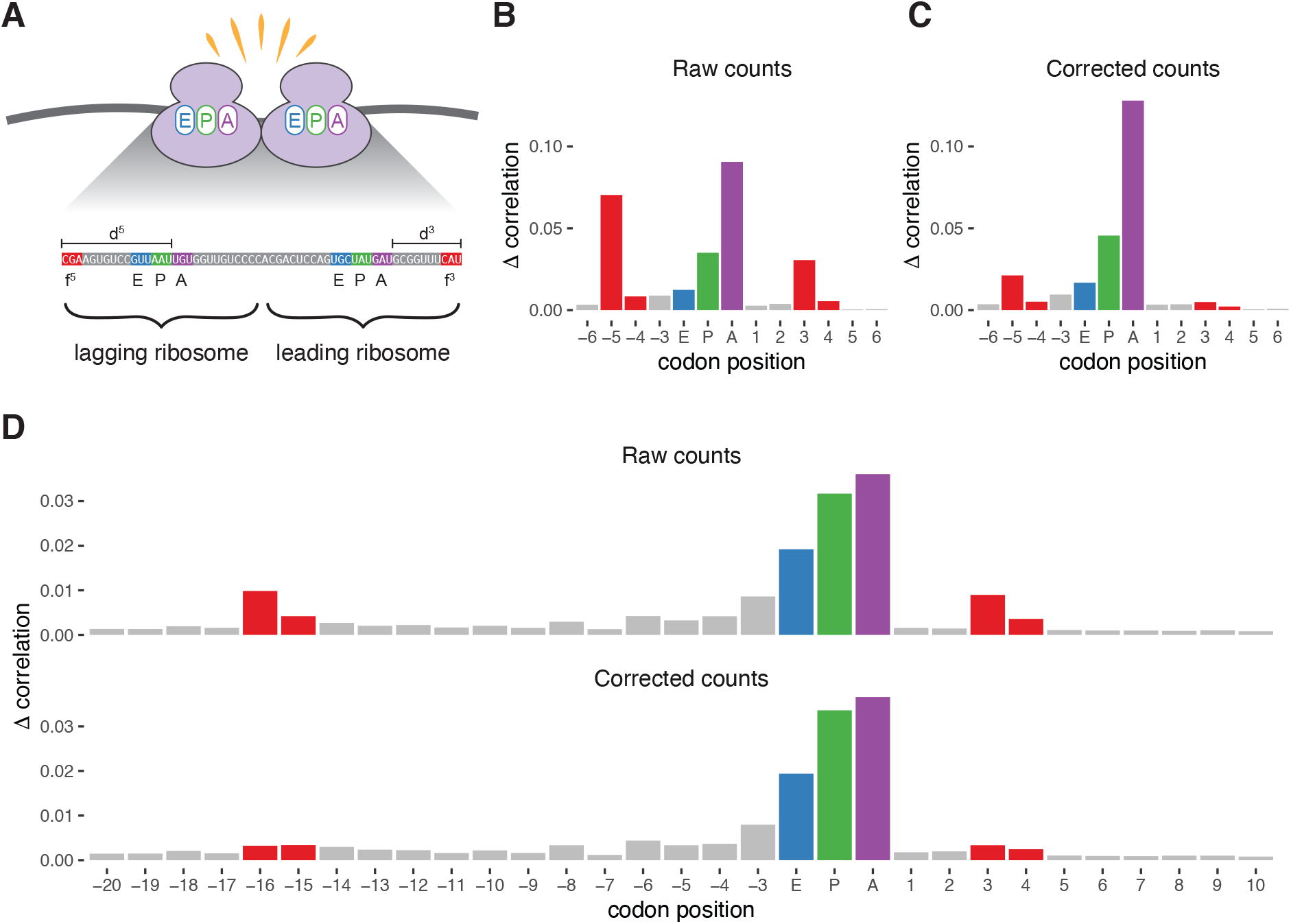
Bias correction of disome footprint counts with correction factors learned from monosome counts. **(A)** Schematic for the featurization of disomes footprints. Rules for assignment of the leading and lagging A-site codon positions are established by metagene plots of footprint counts near start and stop codons. Enumeration of the 5′ and 3′ digestion lengths and bias sequences enables bias correction with `choros`. **(B)** Position importance plot for raw monosome counts. **(C)** Position importance plot for bias-corrected monosome counts. The bias correction factors calculated from monosome data were then applied to disome footprint counts. **(D)** Position importance plot for raw and bias-corrected disome counts. The purple, blue, and green bars correspond to the A-, P-, and E-site codons of the leading (3′) ribosome, respectively.

We applied this approach to a dataset of monosome- and disome-protected fragments [30]. Position importance plots generated for monosome footprints demonstrated the canonical *−*5 and +3 signals indicative of the presence of ligation biases, which were attenuated after performing bias correction with `choros` (Figure 7B-C). We next approached applying monosome bias correction factors to disome data. Because these correction factors are specific to digestion lengths as well as bias sequences, we needed to identify the A-site codon positions of both the leading and lagging ribosome within the disome fragment in order to correctly determine the 5′ and 3′ digestion lengths (Figure 7A). A-site assignment of the 5′ lagging ribosome was performed as in monosome data, where A-site offsets were individually generated per read length and 5′ end framing based on start codon metagene plots. A-site assignment of the 3′ leading ribosome used a similar approach, where read length- and frame-specific offsets were generated based on stop codon metagene plots (Supplementary Figure 6). We then performed bias correction on disome counts using bias correction factors estimated from monosome counts.

To examine whether bias correction with `choros` was able to remove sequence artifacts in disome counts, we generated position importance plots across the span of a disome footprint. We observed an increase in signal at the positions corresponding to the disome fragment boundaries in raw counts, indicative of sequence artifacts as also seen in monosome data (Figure 7D).This signal was dampened in corrected counts, demonstrating that bias correction factors generated from monosome data can indeed be transferred to disome data to correct ligation biases common to both datasets. We recommend that monosome profiling be conducted when collecting footprints of two or more ribosomes so that sequence-based biases may be estimated and properly corrected.

## Discussion

Ribosome profiling informs our knowledge of gene expression by quantifying translation of mRNAs. Beyond this, it also provides a codon-resolution snapshot of ribosome distribution across transcripts, allowing inference of the decoding time of different codons and providing insight into molecular mechanisms of translation. This codon-level information has led to advances in synthetic mRNA design and discovery of amino acid limitations and tRNA alterations in cancers [13, 46–48]. However, faithful recovery of ribosome positions by ribosome profiling presents substantial challenges.

Sequence-dependent biases are pervasive across ribosome profiling datasets, even when careful attention is paid to the library preparation protocol. This arises from the necessary biochemical manipulation of ribosome-protected footprints to make them amenable to high throughput sequencing. For instance, most library preparation approaches require ligation of sequencing adapters onto the ends of the footprints. Ligases generally have sequence preferences, likely for both the 5′ and 3′ entities to be ligated, making these biases unlikely to be completely avoidable. The resulting over- and under-representation of certain footprints leads to distorted ribosome distributions, which can be detected as artifactual signals showing an influence of several nucleotides at either end of footprints on footprint recovery [13, 15, 16]. New ligation-free methods such as the template jumping approach used in OTTR ribosome profiling greatly ameliorate sequence preferences, but cannot completely avoid them [49, 50]. Removing these sequence signals is thus necessary to reveal accurate and precise patterns from ribosome profiling data and to uncover mechanisms of translation regulation and their biological functions.

Our method, `choros`, directly estimates ligase preferences and generates correction factors to produce footprint counts that are free from these sequence-dependent biases. Careful statistical analysis, use of simulated data, and comparisons with our experimental data on ligation preferences confirm the validity of our model. `choros` was able to reverse the effects of ligation biases across many experimental datasets, including disome footprints, and also simultaneously produced parameter estimates that were interpretable and comparatively free of sequence biases. Using our correction method, we showed that a signal often interpreted as a biological phenomenon was potentially the result of biased footprint recovery. Our method will therefore allow more accurate detection of biologically relevant ribosome pauses: interactions with regulatory proteins; intrinsic pauses due to tRNA abundance, peptide bond formation, or nascent chain interactions; collisions between ribosomes; and other as-yet-undiscovered phenomena.

The quantitative model stems from a conceptual data-generating process that can be generalized to multiple library preparation protocols, including dual ligation and template jumping approaches. However, even with `choros`-corrected footprint counts, interpretation of codon-level data from ribosome profiling presents challenges. Ribosome profiling datasets can also suffer from other types of biases. For instance, translation elongation inhibitors such as cycloheximide, used to stabilize ribosomes on mRNAs, can lead to ribosome accumulation on certain codons [51, 52].The data-generating process currently assumed by `choros` does not account for any distortions of ribosome distribution that might occur prior to nuclease digestion.

The sparsity of footprint data on all but the most abundant or highly translated transcripts presents a known limitation in interpreting ribosome profiling data. This limitation extends to our bias correction method. In fact, when no footprints are recovered at a position, whether because of sparse ribosome density or poor footprint recover, bias correction is not possible; `choros` adjusts footprint counts in a multiplicative manner that cannot add footprint counts to positions where none were originally observed. In this analysis, we have used only the most highly translated transcripts to learn the biological and technical parameters governing ribosome distribution. We emphasize that the combination of sparse data and sequence biases renders many transcripts unsuitable for detailed individual analysis by ribosome profiling, while aggregate analyses of patterns are highly reliable. As such, we consider `choros` most appropriate for analyzing translation of abundant transcripts in high-coverage datasets. To mitigate the impact of missing data, future work could explore the addition of pseudocounts in order to restore counts to positions where biased recovery led to no observations, and to increase the reliability of ribosome distributions on sparse transcripts.

In summary, `choros` provides a much-needed solution to the problem of sequence-based biases in ribosome profiling data and should be incorporated into standard analysis pipelines to provide faithful codon-resolution measurements of translation and enable biological discovery.

## Acknowledgements

We thank Premal Shah and the riboviz team for insight and discussion throughout this project, and Nicholas Ingolia, Lucas Ferguson, and Kathleen Collins for discussion and access to unpublished ribosome profiling datasets.

## Funding

This work was supported by the U.S. National Science Foundation [1936069 to L.F.L.], the National Institutes of Health [National Cancer Institute R21CA202960 and National Institute of General Medical Sciences R01GM132104 to L.F.L.], the U.K. Biotechnology and Biological Sciences Research Council [BB/S018506/1 to E.W.J.W.], and the Wellcome Trust [208779/Z/17/Z to E.W.J.W.]. R.T. was supported by the U.S. Department of Defense through the National Defense Science & Engineering Graduate (NDSEG) Fellowship Program.

---

**Supplementary Figure 1.**
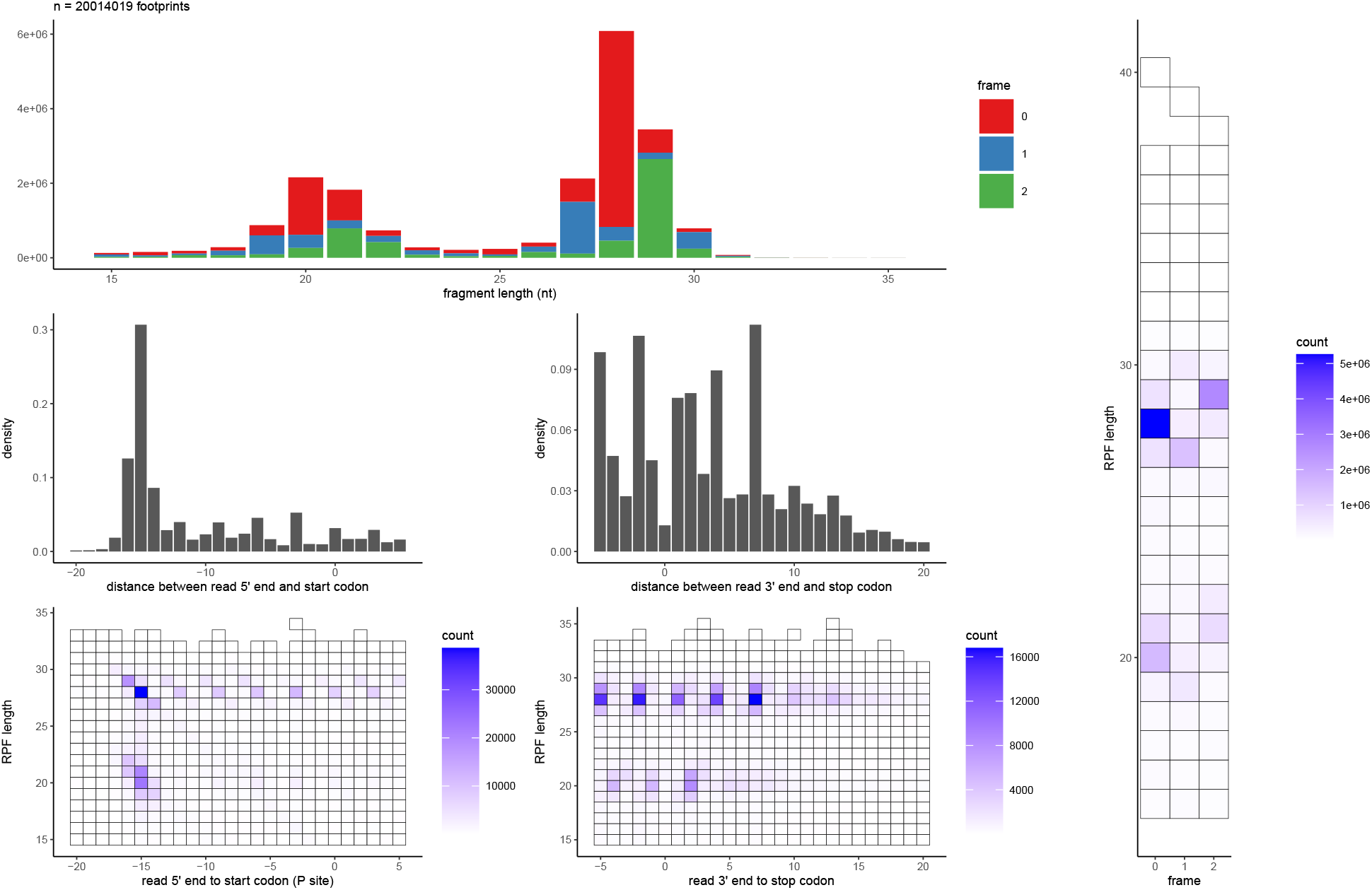
Establishing A-site offset rules from a start codon metagene plot. This plot reflects data generated by Tunney et al. (2018) [13].

**Supplementary Figure 2.**
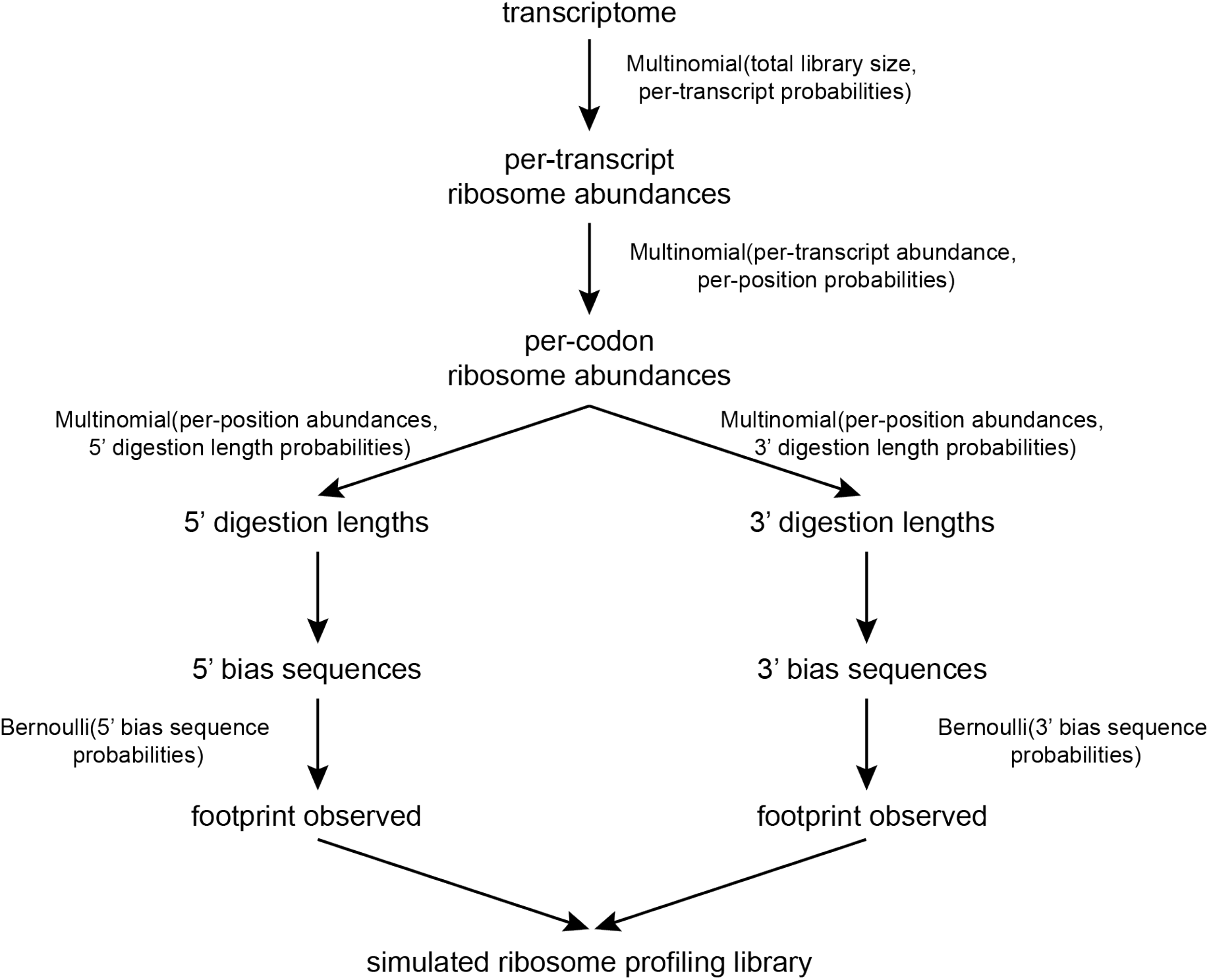
Schematic diagram for the generation of simulated ribosome profiling datasets using `simRiboSeq`.

**Supplementary Figure 3.**
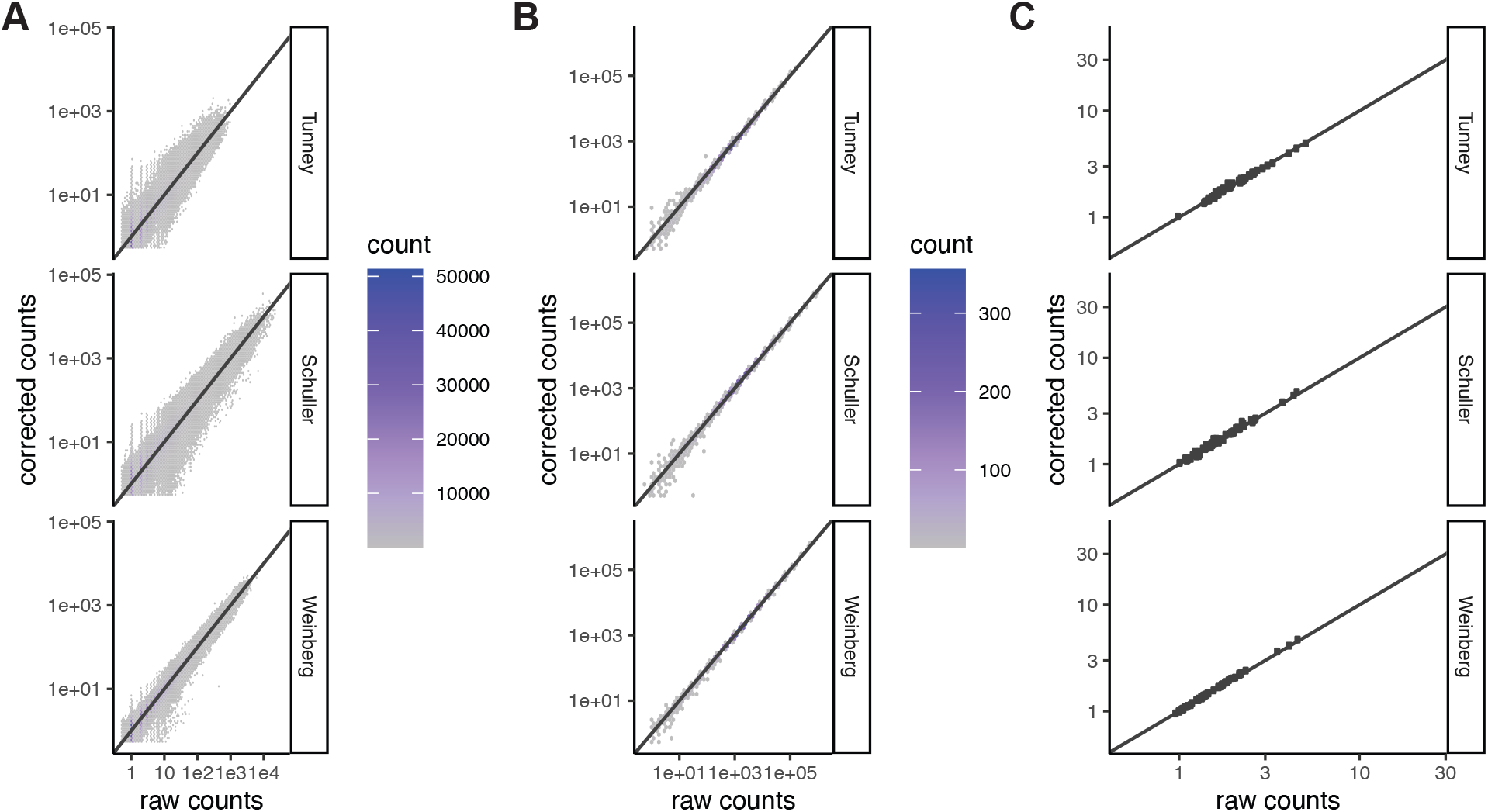
Impact of bias correction on ribosome abundances. **(A)** Per-codon footprint counts, before and after bias correction. Footprint counts were summed by codon position. **(B)** Per-transcript footprint counts, before and after bias correction. **(C)** Codon pause scores, before and after bias correction. Codon pause scores were calculated as follows: footprint counts were aggregated by codon position and normalized to the mean read coverage by transcript (omitting the first and last 20 codons). Normalized footprint counts were averaged by codon identity across the transcriptome.

**Supplementary Figure 4.**
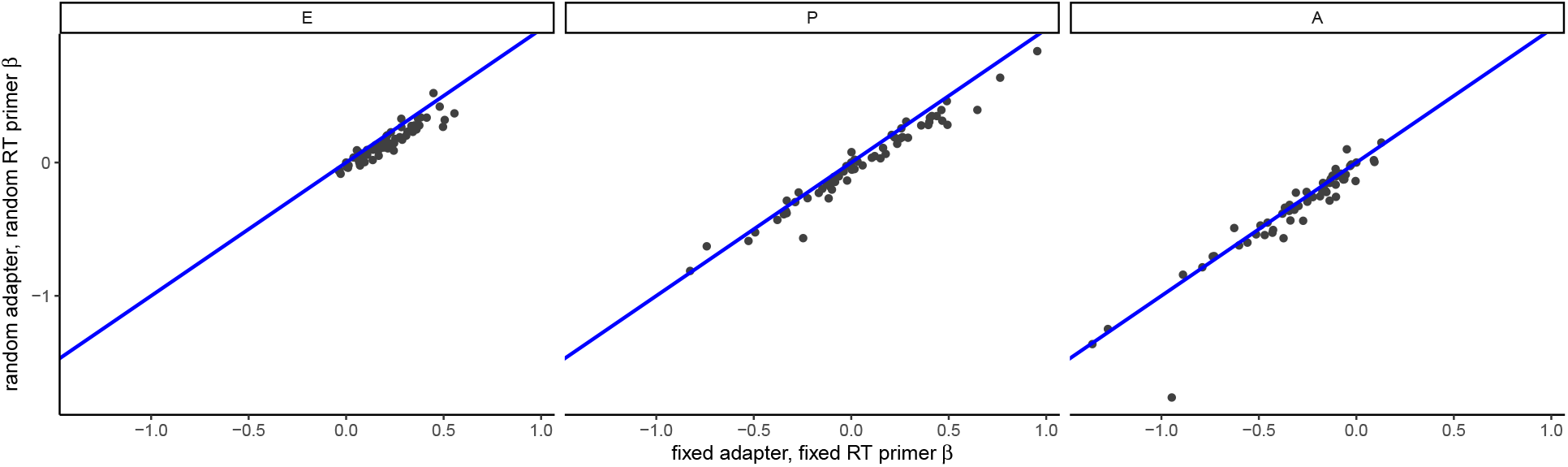
Comparison of codon-level `choros` regression coefficients in an experimental validation dataset. Comparison of A-, P-, and E-site regression coefficients between two datasets generated from the same pool of ribosome footprints but using different library preparation protocols. A 1:1 line is shown in blue.

**Supplementary Figure 5.**
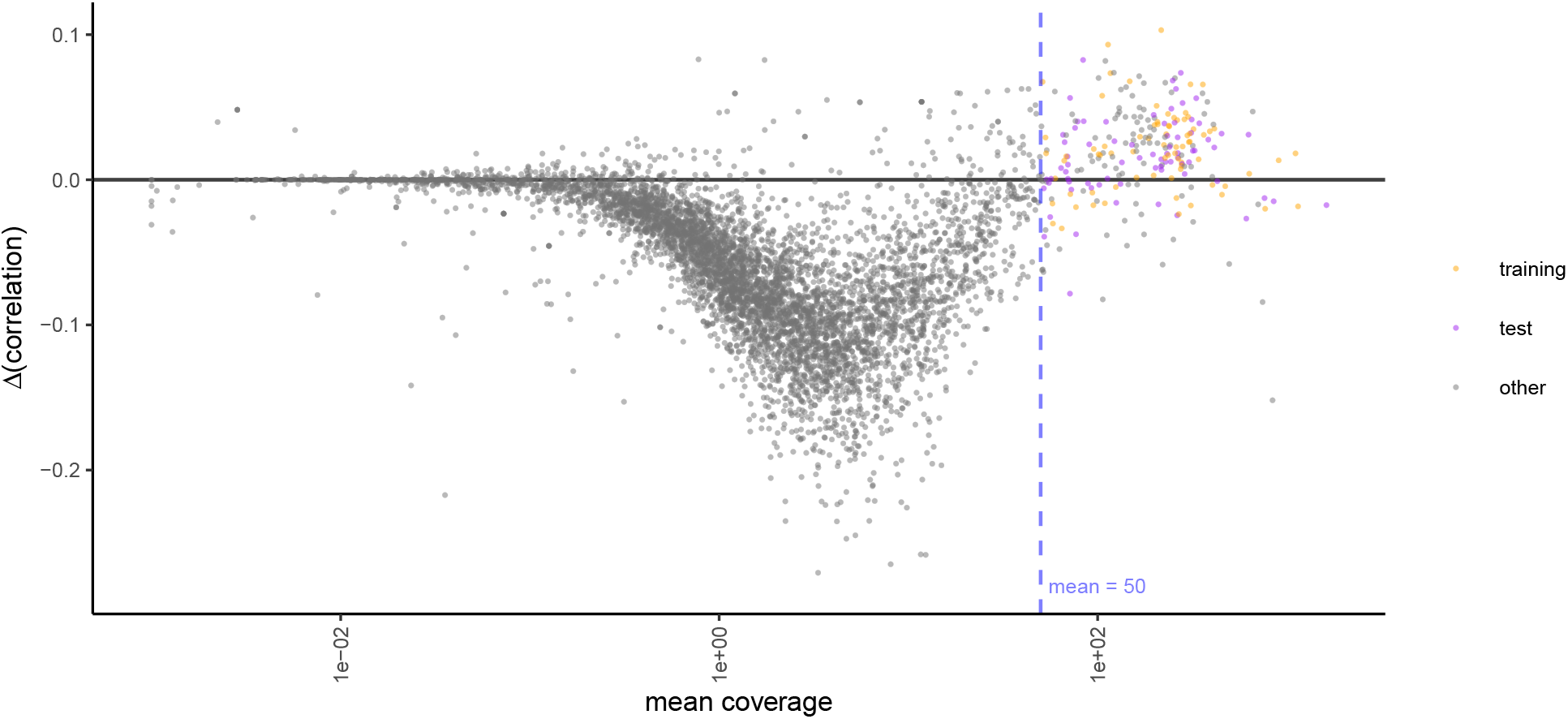
Per-codon footprint counts on sparse transcripts may be untrustworthy. Correlation of footprint counts between library preparation protocols were computed on a per-transcript basis using raw and bias-corrected footprint counts. The y-axis reflects the difference in correlation between protocols when using bias-corrected footprint counts. Only transcripts with a mean coverage of *>*50 footprint counts per codon position tend to increase in similarity after bias correction.

**Supplementary Figure 6.**
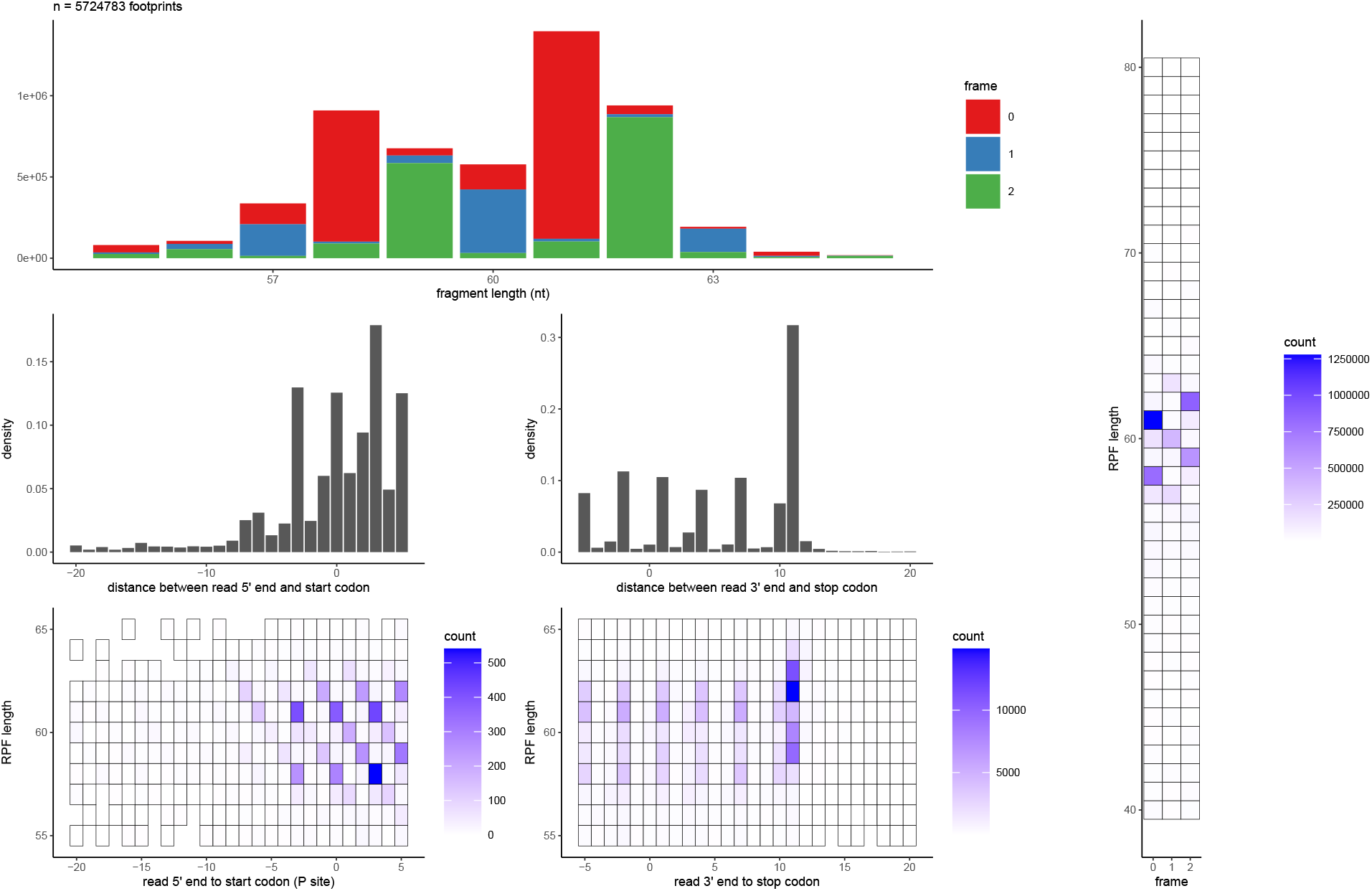
Establishing leading and lagging A-site offset rules for disomes using start and stop codon metagene plots.

## Notes

### Competing Interest Statement

The authors have declared no competing interest.

